# The microbiota of a wild butterfly species over three decades of climate change

**DOI:** 10.64898/2026.06.08.730914

**Authors:** Linyang Sun, Natalie E. van Dis, Andréa Davrinche, Marjo Saastamoinen, Johan Ekroos, Anne Duplouy

## Abstract

As microbes can support essential functions related to many aspects of host physiology, changes in microbial communities can affect host physiology and fitness. Although the effects of thermal conditions on host-associated microbiomes have been demonstrated in laboratory settings, how climate change might affect the structure and functionality of microbial communities in wild populations remains poorly understood. We took advantage of the long-term field survey of the Glanville fritillary butterfly (*Melitaea cinxia*) metapopulation on the Åland Islands, in the Baltic Sea, to fill this gap. We investigated whether bacterial communities associated with larvae showed gradual temporal change in response to slow environmental warming across a 28-year period, or whether these communities responded through abrupt change following a sudden drought event that triggered bottlenecks in their butterfly host population. Using 16S rRNA metabarcoding and metagenomic sequencing, we first showed that *M. cinxia* harbours a set of stable resident bacteria, including *Pseudomonas*, *Telluria,* and Enterobacteriaceae bacteria. We also characterized a gradual shift in the *M. cinxia* associated bacterial community over three decades of increasing temperatures and decreasing precipitations. This shift was not unidirectional for all bacterial taxa, as the dominant *Telluria* and *Pseudomonas* showed opposing responses to environmental trends. Additionally, the 2018 extreme drought, which triggered acute host population bottlenecks, was associated with a severe disruption of *M. cinxia* microbiota, and the loss of key Enterobacteriaceae taxa. However, the *M. cinxia* bacterial community appeared to recover towards pre-drought structure in subsequent years, suggesting resilience to acute climatic perturbations in this system.

## 1. INTRODUCTION

Global temperatures are projected to reach 2.7°C above preindustrial levels by 2100, and even earlier at the poles [1]. The consequences of global warming already include accelerated biodiversity loss [2, 3], increased selection pressures on local populations [4], and changes in the metabolism, physiology and immunity of diverse organisms [5–7]. Stress from heat can also disrupt host-microbe associations [8, 9], and ultimately lead to the host population extinction [10]. For instance, under warm laboratory conditions, aphid mortality increased due to the loss of their obligate symbiont Buchnera aphidicola, which provides the aphids with essential amino acids absent from the plant-sap diet [11]. Alternatively, some microbes can buffer their hosts against environmental thermal stress. For example, thermal tolerances in *Drosophila melanogaster* [12], *Bactrocera dorsalis* [13], and *Ischnura elegans* [14] significantly improved after transplanting naïve hosts with bacteria from conspecifics reared under higher temperatures. Similarly, under heat stress, aphids carrying facultative symbionts *Regiella* or *Fukatsuia* displayed higher fecundity than uninfected aphids [15].

Beyond the direct effects of heat on the host-associated microbes, climatic changes can also indirectly impact insect microbiomes by altering their environmental microbial pools [16]. Changes in temperature and precipitation can modify host-plant quality and leaf-surface chemistry, shifting the microbial species that insects encounter and potential acquire while feeding on plants [17, 18]. In nature, the direct and indirect effects of climate change on the microbiome of wild species can operate simultaneously, and cumulate over long period. Evaluating such changes under natural settings requires long-term field studies.

Long-term microbiome studies of wild populations are confined to a few species. Decades-long microbiome studies of wild meerkats in South Africa [19], and spotted hyenas in Kenya [20] have revealed the adaption and evolution of host-microbiota relationships in response to slow environmental changes. While Risely et al. (2023) revealed that rising temperatures progressively shifted gut communities towards enriched disease-associated bacteria and depleted beneficial mutualists in meerkat populations [19]. Rojas et al. (2023) described temporal shifts in gut microbial composition of hyenas without any effects on their core microbial species [20]. These different responses may reflect difference in the host species ecology, including gut structure, environment, or diet [21]. Hence, these few mammal studies are unlikely to be representative of other taxa, especially of insects, which can exhibit great population fluctuations over short time periods, and experience more dynamic microbial community shifts.

The long-term survey of the Glanville fritillary butterfly (*Melitaea cinxia*, Linnaeus 1758) metapopulation system in the Åland Islands offers a unique opportunity to investigate three decades of microbiome dynamics in a wild insect. Situated in the Baltic Sea, at a latitude >60°N, the Åland Islands are experiencing faster warming compared to more temperate regions [22], and local temperatures are already on average above 2°C higher than 30 years ago (Fig.1B). This butterfly metapopulation has been systematically surveyed every year over the last three decades (i.e. since 1993) [23, 24], to investigate how population dynamics [25, 26], population genetics [27, 28], genomics [29, 30], and adaptability of *M. cinxia* changed with habitat fragmentation and climate change [31]. To date, the gut microbiota of *M. cinxia* has been described as mostly transient [32], highly variable between and within host families [33], and potentially involved in the host immune response to stress [17, 34]. Yet, it remains unclear whether and how the microbiome has changed over time under the warming climate.

**Figure 1.**
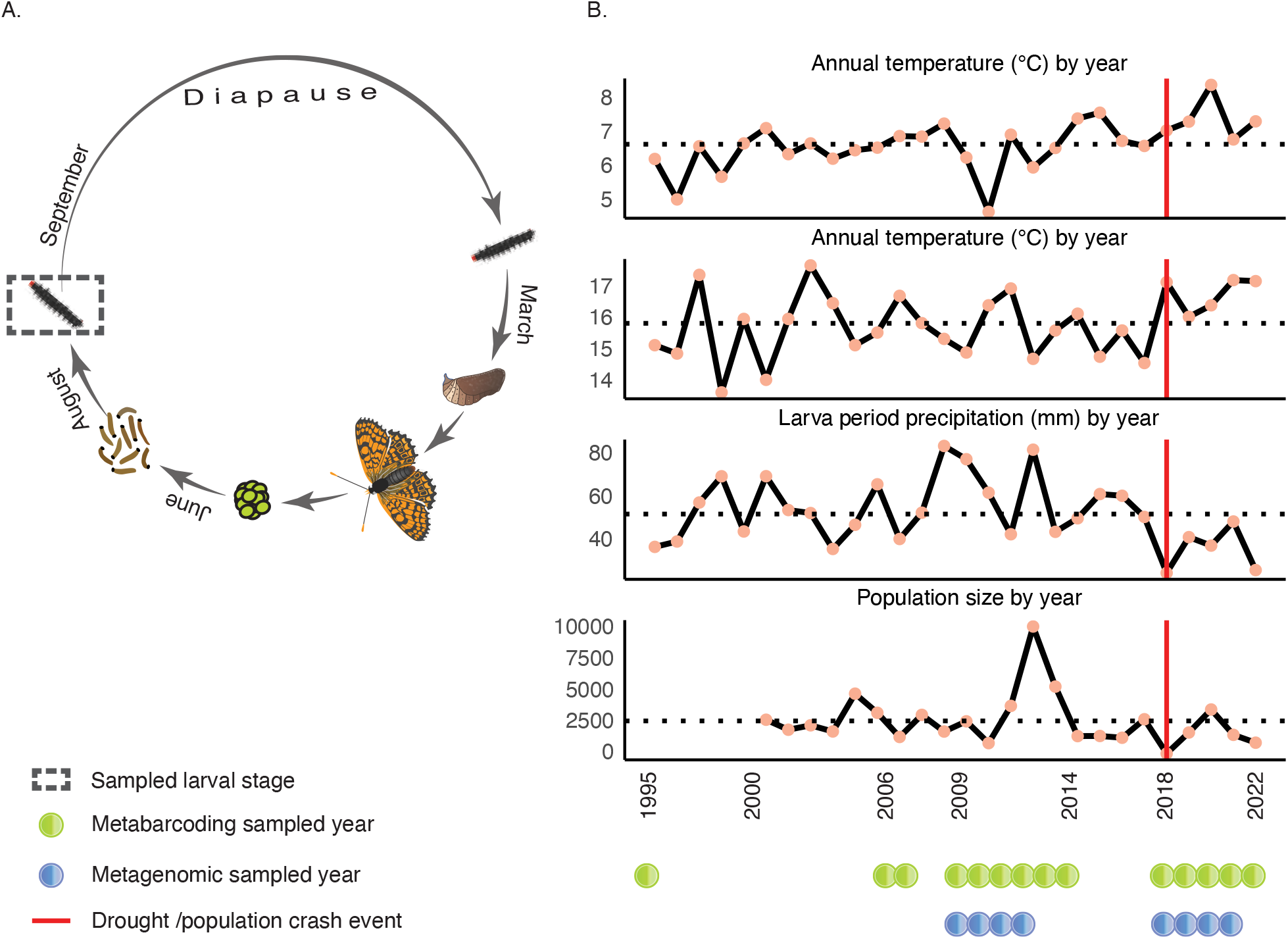
Ecological and environmental background of the *Melitaea cinxia* metapopulation in the Åland Islands, Finland. (A) Lifecycle of *M. cinxia* in the Åland Islands. Diapausing larvae were collected in early September. (B) Fluctuations in the climatic conditions and metapopulation dynamics of *M. cinxia* in the Åland Islands over a 28-year period (1995-2022). Metapopulation data before 2000 are limited, as only a portion of the populations were surveyed during those early years. We processed larvae from 14 years across the 28-year period through 16S metabarcoding (green dots), and from eight years through metagenomics (blue dots).

To assess long-term changes in the microbiota associated with this wild species, we leveraged the wealth of samples collected across a 28-year period (1995-2022) in the Åland islands, to generate microbial community data through metabarcoding and metagenomic sequencing approaches. We hypothesized that gradual warming associates with directional microbiome turnover across decades, whereas extreme climatic events produce abrupt community shifts. By linking long-term microbiome dynamics with climatic variation, we test whether warming and extreme weather have left detectable signatures in the microbiota of a natural insect population.

## 2. MATERIAL AND METHODS

### 2.1 Study system

*Melitaea cinxia* occurs across temperate regions of Europe, North Africa and Asia[23]. In Finland, the species is restricted to a network of meadows and pastures scattered across the Åland Islands[35]. Adults mate from June to early July, and do not disperse further than a couple of kilometres away from their oviposition sites[23]. Females lay clutches of >50eggs on two host-plant species: *Plantago lanceolata* (Linnaeus, 1753) and *Veronica spicata* (Linnaeus, 1753)[36]. After hatching, larvae live communally in silk nests throughout most of larval development, including overwintering diapause from September until pupation in April [37, 38] (Figure 1A).

Since 1993, larval silk nests have been surveyed annually across the Åland metapopulation [24]. In the autumn of each year, field assistants have systematically screened up to 4 400 geolocated meadow habitats, and recorded all detected larval silk nests [24]. Larvae have been collected from the observed nests, labelled, and preserved at -20°C [24].

To analyse changes in the composition of the bacterial community associated with *M. cinxia*, we selected 570 larvae, collected on 14 years between 1995 and 2022, for metabarcoding sequencing (Figure 1B) (see section 2.3). Additionally, to further characterize changes in the genomics and functionality of key microbes associated with *M. cinxia*, we screened an additional 298 *M. cinxia* whole genome resequencing projects of larvae collected during 2009–2012 and 2018–2021 (Figure 1B) (N.E. van Dis, unpubl. data) (see section 2.3). To reduce the potential family genetic effect, for each year, we only selected larvae found on different sites. To reduce the effect of diet on the microbiota [33], we only selected larvae collected on *P. lanceolata*. Out of the 298 specimens used for the whole genome resequencing project of *M. cinxia*, 43 were also used in the metabarcoding sequencing (see metadata in Supplementary File 1).

### 2.2 Climatic data

To examine how climatic conditions might influence microbial communities associated with *M. cinxia* larvae, we retrieved meteorological data for the years 1995 to 2022 from the Finnish Meteorological Institute’s open data portal (https://en.ilmatieteenlaitos.fi/). We used data from the Jomala weather station, which is central to the Åland Islands. These station-level measurements do not capture microclimatic variation across sampling sites but rather provide Island broad spatial resolution. From this data, we calculated annual mean temperature, annual mean precipitation, and mean temperature and mean precipitation during the larval period only (i.e. between June 1^st^ and August 30^th^ of each year) (Figure 1). After testing for collinearity by calculating Variance Inflation Factor (VIF) <10 by using R package “car”[39], we only kept three of these climatic variables: annual mean temperature, larval period temperature, and larval period precipitation.

### 2.3 Library preparation and sequencing

For the 16S metabarcoding approach, we used DNA extracts previously generated from other studies (N=474), and DNA extracts (N=96) produced specifically for this study. All DNA extracts were produced using the Qiagen DNeasy Blood and Tissue extraction kit (Qiagen, Germany), following the manufacturer’s protocol. We then amplify the V5-V6 hypervariable region of the 16S rRNA gene, using the 784F/1061R primers [40]. All samples were evenly distributed in six 96xwell plates, each including a sterile water control. All samples were amplified in duplicate (Supplementary File 1) and pooled for sequencing through a 300-paired-end run on a MiSeq v3 Illumina Sequencing platform at the Finnish Institute for Molecular Medicine (FIMM). For the metagenomic approach, the DNA was extracted for the original purpose of another study (N.E. van Dis, unpubl. data), using three different methods (see Supplementary file 1). DNA extracts were sequenced in three batches on the NovaSeq Illumina (PE150) platform at BGI Genomics and Novogene Europe, with a median sequencing coverage of 25–37X.

### 2.4 Bioinformatic and statistical analysis

We conducted data curation and analyses using the Puhti server facility and bioinformatic tools provided by CSC (IT Center for Science, Finland). All statistical analyses were conducted using R version 4.4.2 [41], with the *vegan* and *compositions* packages. The different R packages used for the analyses are described briefly below, and in more details in the supplementary file 1.

#### 2.4.1. Metabarcoding data

We used the QIIME2 platform [42] to conduct primer removal, quality filtering, DADA2 denoising, and taxonomy assignment using Greengenes2 [43]. Sequences amplified from negative controls were subtracted from the biological samples to control for PCR contamination [34]. We excluded all amplicon sequence variants (ASVs) with total read counts <10, and all samples containing <20 unique ASVs. The dataset was rarefied to 1,000 reads per sample based on the sequencing depth distribution (Figure S2). After these quality checks, the dataset included N=396 specimens (70%, Figure S1). A As plant-associated bacteria can significantly influence the microbial community composition of *M. cinxia* larvae [33], we isolated the plant microbes from our data following the results of [33]. We trained SourceTracker2 [44] to estimate the percentage of plant bacterial taxa in our datasets. We then applied a differential abundance analysis (DESeq2) [45] to identify taxa (at genus level) enriched only in plants in the reference datasets, and removed them from our dataset.

To assess changes in the alpha-diversity of the bacterial community associated with *M. cinxia*, we first calculated the Shannon diversity and Observed species richness indices using the phyloseq package [46]. The two indices inform on community richness and evenness. The relationship between these alpha-diversity indices and year was assessed using a linear model.

To reduce noise in the beta-diversity analyses [47], due to large number of rare bacterial species in our datasets, we applied a Multivariate Cutoff Level Analysis (MultiCoLA) [48]. The MultiCoLA allowed us to evaluate the contribution of each ASV to overall community variation. Based on this contribution, we split the community into “rare” and “abundant” subgroups. We used distance-based redundancy analyses (db-RDA) to test whether year and climatic variables significantly explained variation in the composition of *M. cinxia*‘s whole, abundant, and rare bacterial communities. These models were evaluated using Bray-Curtis, Jaccard, and weighted UniFrac dissimilarity matrices (capscale, Vegan package) [49]. The significance of all models was determined using permutational multivariate analysis of variance (PERMANOVA; with 999 permutations) [50]. We fitted the variables year, annual temperature, larval-period temperature, and larval-period precipitation in the models. By projecting species scores onto the first two constrained axes, and ranking genera by their respective Euclidean distances from the origin, we determined the weight of the different bacteria on the community composition. We also built zero-inflated negative binomial distributions within Generalized Linear Latent Variable Models (GLLVMs) using the package ”gllvm” [51] to quantitatively assess the effect of climate on each taxonomic level. We selected larval-period precipitation as an explanatory variable for all GLLVMs, as it emerged as the dominant driver of beta-diversity in the db-RDA. Given the potential for high inter-individual variation in *M. cinxia* microbial communities [33], we also included specimen IDs as a random factor.

#### 2.4.2. Metagenomics

We use bwa-mem2 v.2.2.1 [52] to remove host reads from the *M. cinxia* whole-genome re-sequencing data. We then controlled the quality of the remaining unmapped sequences using BBTools [53] and fastp [54]. High-quality bacterial reads were assembled into contigs using MEGAHIT [55], followed by metagenomic binning with MetaBAT2 [56], and refinement with MAGpurify [57] to remove incorrectly clustered contigs. We evaluated the quality of each bin using CheckM2 [58], retaining only bins with ≥70% completeness and <5% contamination. The refined bin sets were dereplicated using dRep [59], yielding 25 high-quality Metagenome-Assembled Genomes (MAGs).

All MAGs were assigned taxonomy using GTDB-Tk [60], and functional annotations using Prokka [61]. To investigate phylogenetic relationships between the 25 refined MAGs, we reconstructed a maximum-likelihood phylogeny based on the concatenated alignments of the 25 MAGs to conserved single-copy marker genes in the Genome Taxonomy Database. We trimmed poorly aligned regions using trimAl [62]. The final phylogeny was inferred using IQ-TREE 2 [63] with ModelFinder Plus and 1000 ultrafast bootstrap replicates as parameters, including the genome of *Bacteroides thetaiotaomicron* (NCBI accession #GCF_014131755.1) as an outgroup. We inferred the functional potential of each MAG by evaluating the proportion of enzymatic steps present in each genomic assembly for each KEGG metabolic pathway using KEGGDecoder [64] and dbCAN [65], respectively.

To assess how the community of MAGs changed between 2009-2012 and 2018-2021, we calculated the prevalence and abundance of each MAG bin based on coverage information recovered with CoverM [66]. Per sample bin abundance was quantified as transcripts per million (TPM). To assess the relationship between metagenomic bin composition and selected variables, we performed distance-based redundancy analysis (db-RDA). Zero values were replaced with half the minimum observed non-zero TPM value, and the data were centered log-ratio (CLR)-transformed to account for the compositional nature of the data. Euclidean distance on CLR-transformed values was used as a dissimilarity measure equivalent to Aitchison distance. We fitted the db-RDA model using the capscale function in the R package *vegan*, with year, annual mean temperature, larval-period mean temperature and mean precipitation, as constraining variables. Statistical significances of the overall model and of the individual terms were assessed using permutation-based ANOVA (999 permutations). Species scores were assigned to identify the metagenomic bins most strongly associated with the constrained axes.

Finally, we evaluated whether the MAGs abundances and prevalences significantly varied between the pre-drought years (2009–2012), the drought year 2018, the post-drought year 2019, and later years (2020–2021). Given the unequal sample sizes across these periods, non-parametric methods were employed for this analyses using the dunn.test package [67]. For abundances, we performed Kruskal-Wallis tests on log-transformed TPM (log(TPM+1)) for each MAG, followed by Dunn’s post-hoc pairwise comparisons with Benjamini-Hochberg correction. For prevalence, we used chi-squared tests for each MAG, followed by pairwise Fisher’s exact tests with Benjamini-Hochberg correction.

## 3. RESULTS

### 3.1 The microbial community associated with *M. cinxia*

Based on SourceTracker2 sink-source prediction results, we found that more than one third of the bacteria associated with diapausing *M. cinxia* larvae are likely from plant (Figure S1; Table S1). The DESeq2 differential abundance analysis (p < 0.05, log_2_FoldChange>3) identified 25 genera, predominantly *Sphingomonas, Methylobacterium, Variovorax, Mycobacterium, and Xanthomonas*, as enriched only in the plant (Table S2, Figure S3). After removing this contamination from the plant, 880 bacterial genera remained.

Each larva included 20 to 112 bacterial genera (median=41; Figure S4), and both alpha-diversity indices (i.e. Shannon and richness) remained similar across years (Figure 2B; Table S11). The microbial community associated with *M. cinxia* larvae exhibited a core-dominated structure characterized by few dominant genera, in addition to numerous rare taxa. Two genera, *Pseudomonas* (family Pseudomonadaceae) and *Telluria* (family Burkholderiaceae), dominated the community with the highest mean relative abundances (0.17 and 0.14, respectively), and high prevalences (>80%) across all years (Figure 2A; Table S4). These two genera were shared by the host plant and larvae reference databases [33](Table S2). In contrast, *Acinetobacter* (Moraxellaceae), *Corynebacterium* (Mycobacteriaceae), and *Staphylococcus* (Staphylococcaceae), were highly prevalent (68-77%), but at low relative abundance in each larva (0.024-0.027, Figure 2A; Table S4).

**Figure 2.**
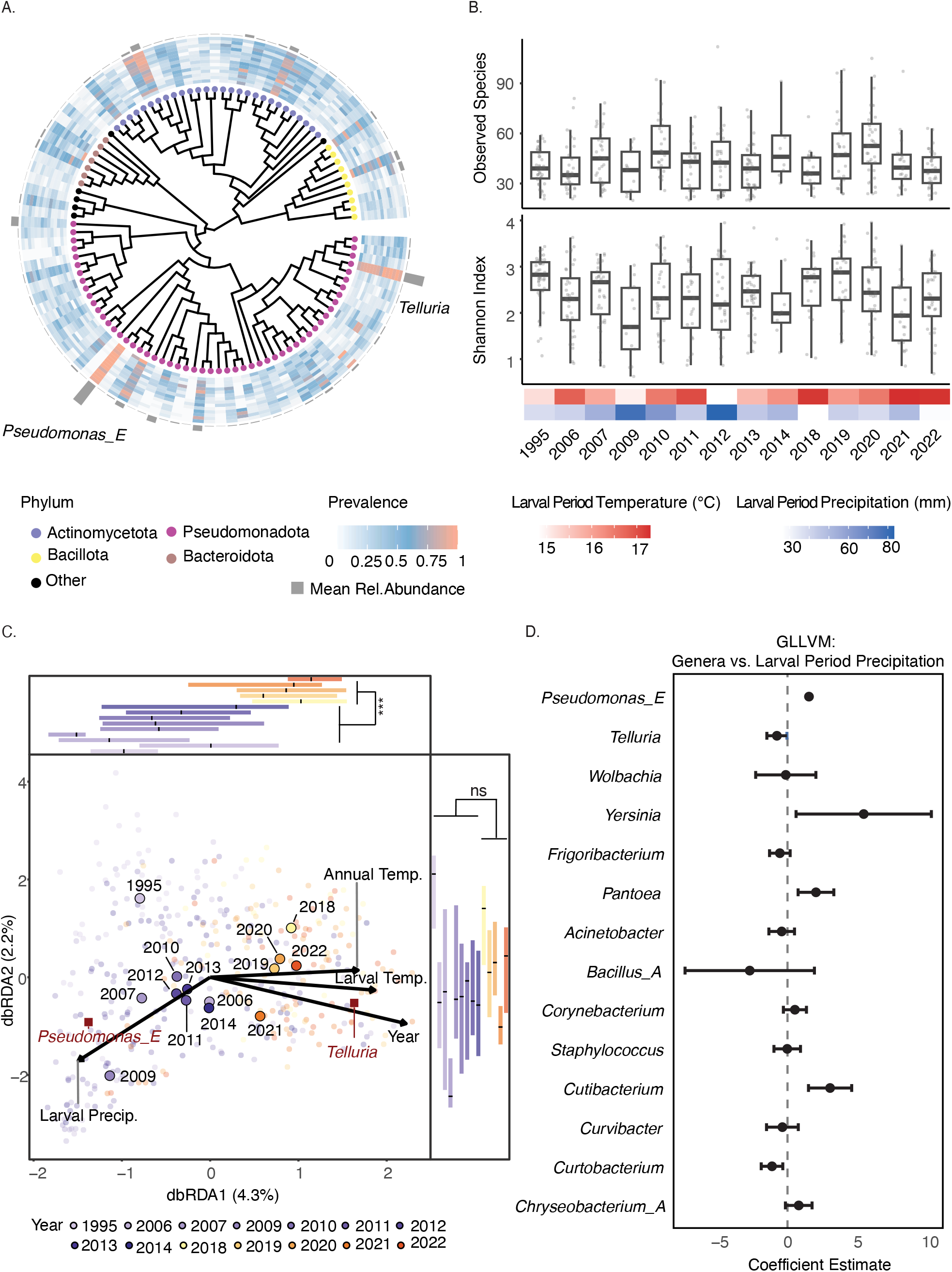
Bacterial community associated with the diapausing larvae of *M. cinxia* and changes observed through years under different climatic conditions. **(A)** Phylo-heatmap of all bacterial genera with a relative abundance greater than 0.1%. The shape of each node at the tree tips indicates phylum information. The heatmap shows the prevalence of each genus for each year from 1995 to 2022 (inner to outer). The outer bar plot shows the mean relative abundance for each taxa, including the two most abundant genera, *Telluria* and *Pseudomomnas*. **(B)** Boxplots of *M. cinxia* bacterial alpha-diversity distribution, as shown by the total number of observed species (top) and the Shannon index (bottom) for each year. The bottom heatmap shows yearly climatic conditions. **(C)** Distance-based redundancy analysis (db-RDA) of weighted UniFrac dissimilarity, illustrating the relationship between microbiome composition and climate variables. The direction of the variables ‘Annual Mean Temperature’, ‘Larval Period Temperature’, and ‘Larval Period Precipitation’ are represented by vectors. Each specimen is represented by a circle colored by year, with shades of purple indicating pre-drought years and shades of orange post-drought years (or something along those lines). Only two genera with importance score > 1 are displayed by red squares. The side boxplots show yearly distribution of specimens across each db-RDA axis. **(D)** GLLVM coefficient estimates of the top 15 larval genera against larval period precipitation. Positive/negative values reflect increasing/decreasing abundance with precipitation, respectively.

By screening metagenomic data from *M. cinxia* re-sequencing projects (vanDis, unpubl. data), we recovered 25 Metagenome-Assembled Genomes with highest quality (out of 126 MAGs total, Table S8). Among these 25 MAGs, ten belonged to the genus *Pseudomonas* (Pseudomonadaceae) and nine belonged to different genera within Enterobacteriaceae. The remaining six MAGs belonged to other families (Figure 3A).

**Figure 3.**
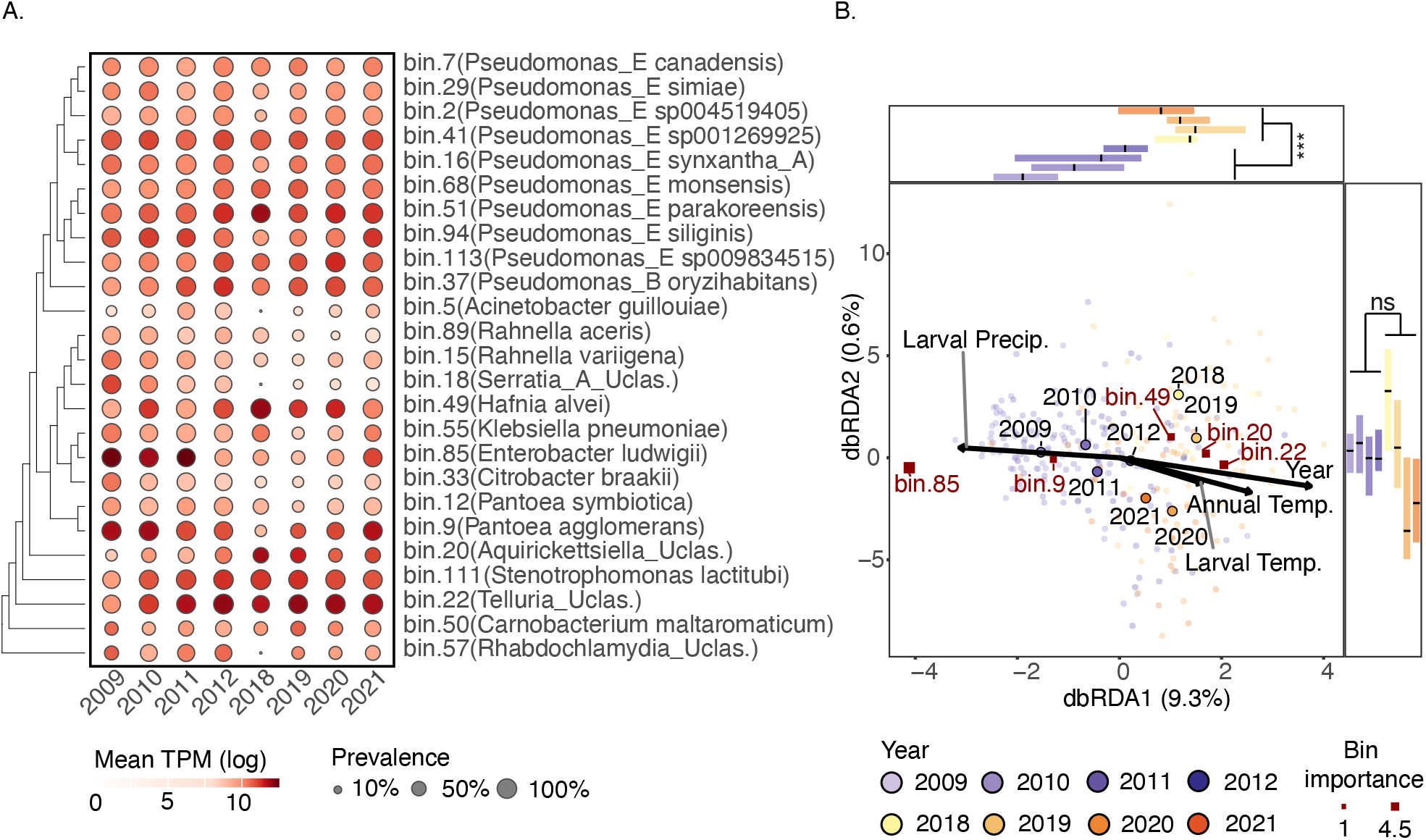
Temporal dynamics, environmental associations, and functional potential of metagenome-assembled genomes (MAGs). **(A)** Mean abundance (log TPM, color intensity) and prevalence (bubble size) of 25 MAGs across sampling years (2009–2021). MAGs are clustered by phylogenetic relationships. **(B)** Distance-based redundancy analysis (db-RDA) of Aitchison dissimilarity, illustrating the relationship between metagenomic bin composition and climate variables (Annual Mean Temperature, Larval Period Temperature, and Larval Period Precipitation). Specimens are represented by small circles colored by sampling year, with shades of purple…etc see above. The central position of each year were calculated and marked as black delimited circles. The top genera are displayed by red squares, with square size reflecting each genus importance score.

### 3.2 *Melitaea cinxia* microbiota response to climate

We tested whether year and climatic variables significantly explained variation in bacterial community composition (Figure S5-13). As all dissimilarity models yielded broad consistent patterns, we selected the weighted UniFrac model for subsequent analyses. The abundant community (i.e., 54 most abundant genera) displayed the largest share of variation (i.e., 6.5% of total variance on both db-RDA axes, Figure 2C, Figure S5-13), and year accounted for the largest proportion of this variation (R^2^ = 0.037), followed by larval-period precipitation (0.020), annual mean temperature (0.011), and larval-period temperature (0.009). In the marginal tests, however, larval-period precipitation exhibited the strongest independent contribution (0.021), followed by year (0.015), annual mean temperature (0.009), and larval-period temperature (0.009)(all *P*<0.001; Table S5). Year and the two temperatures variables, however, acted in the opposite direction to larval-period precipitation (Figure 2C), in agreement with the observed decline in local precipitation levels and the documented rise of temperatures in this region over the study period [68].

The db-RDA revealed that *Telluria* (importance = 1.96) and *Pseudomonas_E* (importance = 1.92) contributed the most to community variation (Table S5). These two genera dominate the bacterial community across years (Figure 2A-C), indicating that temporal shifts in community composition were primarily driven by fluctuations in the prevalent taxa rather than by turnover of rare species. In the db-RDA ordination space (Figure 2C), the contribution of *Telluria* was positively associated with temperature, whereas the contribution of *Pseudomonas_E* was positively associated with the precipitation. Similarly, the GLLVM models identified larval-period precipitation as positively associated with *Pseudomonas_E* (estimate = 1.53, P < 0.001), but negatively associated with *Telluria* (estimate = −0.76, P = 0.038) (Figure 2D; Table S6; AIC = 47134.81, log-likelihood = −23035.41).

For the metagenomic dataset, the Aitchison-based db-RDA model explained the greatest proportion of variance, with dbRDA1 accounting for 9.3% and dbRDA2 for 0.6% of the total variance (Figure 3B; Table S7). Overall, the patterns of the metagenomic db-RDA were broadly consistent with those observed in the metabarcoding-based db-RDA. The positive association with temperatures of *Telluria* (bin22, importance = 2.08) was conserved. However, while *Pseudomonas* abundance was positively associated with precipitation in the 16S rRNA dataset, individual *Pseudomonas* MAGs showed no consistent responses to precipitation in the metagenomic dataset. In fact, the different *Pseudomonas* MAGs displayed contrasting associations with climatic variables: bin.94, (*Pseudomonas_E siliginis;* importance = 0.64) was positively associated with precipitations, while bin.51 (*Pseudomonas_E parakoreensis*; importance = 0.75) and bin.113 (*Pseudomonas_E* sp009834515; importance = 0.75) were not. Additionally, the metagenomic data revealed that bin.9 (*Pantoea agglomerans*; importance = 1.30), bin.85 (*Enterobacter ludwigii;* importance = 4.15) and bin.18 (*Serratia_A* Uclas. ; importance = 0.84) were positively associated with precipitations, while bin.49 (*Hafnia alvei*; importance = 1.44), bin.20 (*Aquirickettsiella*_Uclas.; importance = 1.71), and bin.111 (*Stenotrophomonas lactitubi*; importance = 0.84) were associated with warmer temperatures.

To assess the effect of the 2018 drought on the individual MAGs, we compared their abundances and prevalences across the pre-drought 2009–2012, 2018 drought, 2019 post-drought, and later years 2020–2021 periods (Table S12, Figure S14-15). The MAGs from *Pantoea agglomerans* (bin.9), *Serratia Uclas.* (bin.18) and *Rhabdochlamydia Uclas.* (bin.57) declined significantly in both abundance and prevalence in the 2018drought year compared to earlier years (Dunn’s test and Fisher’s exact test, pre- vs 2018 p < 0.05). Among these, *Pantoea agglomerans* and *Rhabdochlamydia Uclas.* recovered fully by 2020–2021 (pre- vs post-drought p > 0.05 for both metrics), whereas *Serratia Uclas.* showed no recovery (pre- vs post-drought p < 0.001). Conversely, the abundance of *Hafnia alvei* (bin.49) significantly increased in 2018 drought year compared to the pre-drought period (Dunn’s test, pre- vs 2018 p < 0.01) and decreased back to pre-drought levels from 2019.

### 3.3 Functional profiling

After screening the genetic content of each MAGs, we were able to assign metabolic capabilities from 14 pathway categories to these genomes, with completeness ranging from absent to complete (Figure 4; Figure S16; Table S9).

**Figure 4.**
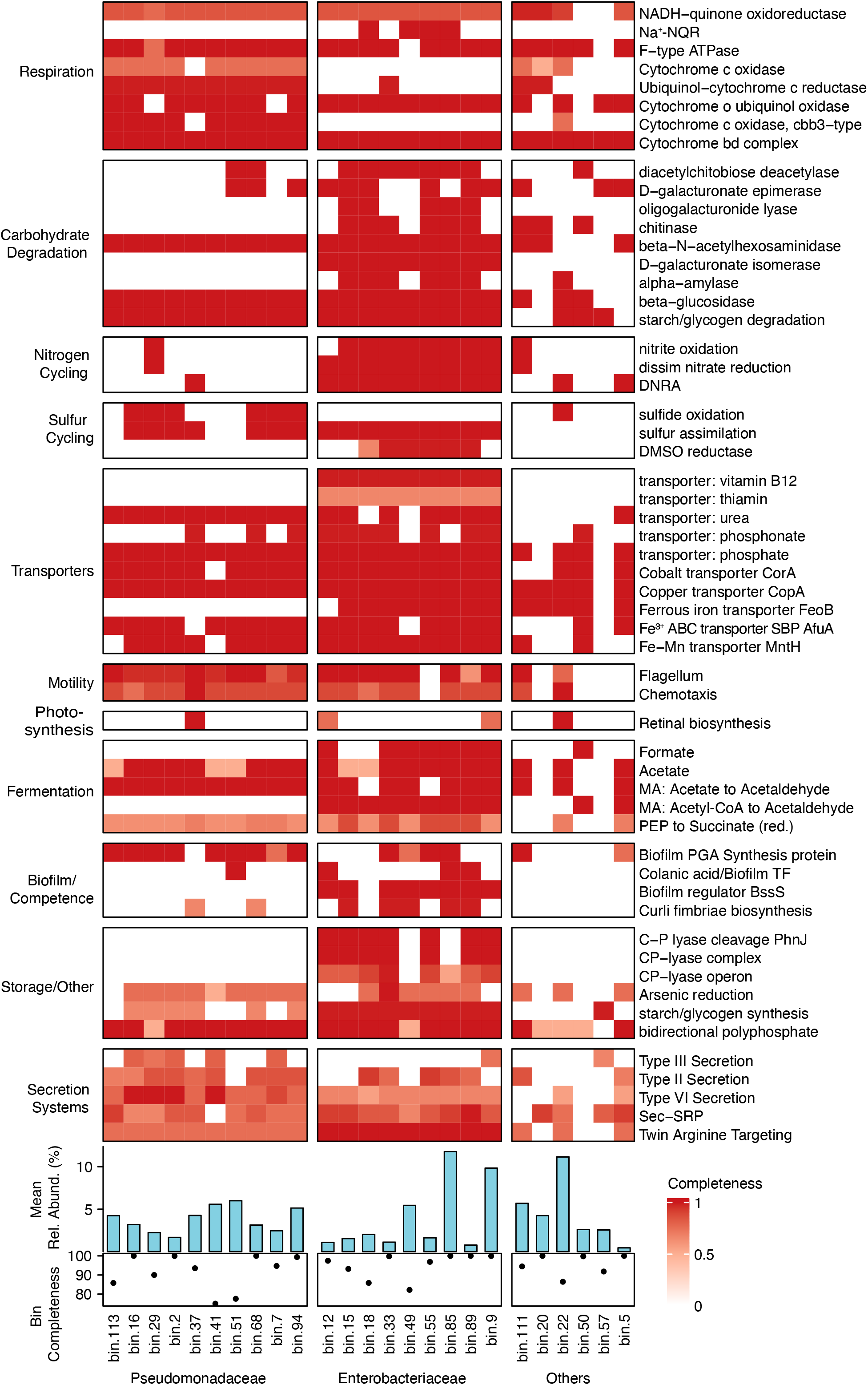
Functional potential of metagenome-assembled genomes (MAGs). Completeness of key metabolic pathway across all 25 MAGs, grouped by taxonomic family. Only differential metabolism pathways are shown in this figure. Side bar plots indicate mean relative abundance (%) and genome bin completeness (%) for each MAG. The illustration was built using heatmaps in ggplot2 [87].

#### 3.3.1 Ten MAGs from the genus *Pseudomonas*

The bacterial genus *Pseudomonas* dominated the *M. cinxia* microbiome across all years and was a key driver of community variation. The ten MAGs from this family comprised heterotrophic, non-photosynthetic bacteria. They shared pathways for central metabolism, aerobic respiration, and motility. These MAGs encoded partial NADH-quinone oxidoreductase and cytochrome c oxidase pathways, and most lacked complete secretion systems. They retained most transporter pathways but lacked transporters for vitamin B12, thiamin, ferrous iron (FeoB), and phosphonate, and some MAGs displayed reduced branched-chain for valine, isoleucine and leucine syntheses. Nitrogen cycling pathways were also largely absent. Their carbohydrate degradation pathways target simple sugars and oligosaccharides (Figure S7).

#### 3.3.2 Nine MAGs from the Enterobacteriaceae family

Enterobacteriaceae was the second most diverse bacterial family from the *M. cinxia* larvae microbiome, and several of its members showed marked declines in prevalence during the 2018 drought. The nine Enterobacteriaceae MAGs span multiple genera, yet they share a broadly conserved metabolic architecture. These MAGs encode heterotrophic, non-photosynthetic lifestyles that are specialized in nitrogen cycling, or sulfur cycling, or both. Their respiratory metabolism is consistently incomplete, suggesting adaptation to low-oxygen conditions in the insect gut. These MAGs can complete aerobic or microaerophilic respiration rather than full oxidative phosphorylation, as they retain the cytochrome *bd* complex, cytochrome *bo* ubiquinol oxidase, and a near-complete F-type ATPase. Most also encode broad carbohydrate degradation capacities targeting plant cell-wall components (Figure S6), suggesting a potential role in host digestion of complex dietary polysaccharides [67]. Complete pathways for nitrogen cycling, central carbon metabolism, amino acid biosynthesis, and substrate transport were universally present across these nine MAGs. Notably, although most of these MAGS are likely from commensal bacteria, four of them (bin.85, 55, 49, and 18) carry the Na - translocating NADH-ubiquinone oxidoreductase, a pathway generally considered pathogen-specific [69].

#### 3.3.3. Six additional MAGs

Of the remaining six MAGs, *Telluria* (bin.22) was mentioned previously, as a dominant member of the *M. cinxia* larval microbiome across all years, while the other five MAGs were rarely detected in the metabarcoding dataset.

(a) The *Telluria* MAG (bin.22), the *Acinetobacter guillouiae* MAG (bin.5), and the *Stenotrophomonas lactitubi* MAG (bin.111) exhibit almost complete amino acid biosynthesis pathways (17, 19 and 17/20 amino acid pathways, respectively), suggesting they do not rely on the host for these essential amino-acids. *Telluria* is the only of the three showing characteristics of a facultative aerobic bacterium, with sulfide oxidation capability, cbb3-type cytochrome c oxidase for microaerobic adaptation, and efficient capacity for aerobic respiration. It is also uniquely motile (chemotaxis and flagellar) and shows retinal biosynthesis for potential rhodopsin-based light sensing.
(b) The *Carnobacterium maltaromaticum* MAG (bin.50) exhibits facultative anaerobic lifestyle with high glycolysis, and possesses unique formate and lactate fermentation, and the highest competence-related components for horizontal gene transfer.
(c) The Unclassified *Rhabdochlamydia* MAG (bin.57) showed the most reduced profile, possibly more characteristic of a symbiotic life-style, with limited amino acid biosynthesis pathways (only glycine, serine, and asparagine), but unique V-type ATPase, Type III secretion system, and starch/glycogen synthesis.

## 4. DISCUSSION

The microbiota of the boreal butterfly population of the *Melitaea cinxia* butterfly harbored a median of 41 bacterial genera, from a general pool of 800 bacterial genera regularly found across the study period. As the Åland population (latitude >60°N) has experienced both gradual and acute climatic changes over the last decades [22], we explored whether the microbial communities associated with M. cinxia larvae changed over a 28-year period (1995–2022). Our results indicated the bacterial community shifted with fluctuations in several climatic variables, and that recovery from those shifts could occur, although with delay. Through functional profiling of bacterial genomes, we further found that out of the taxa sensible to climatic variables, some might encode metabolic pathways relevant for their host fitness. How these changes are likely to affect the future of this butterfly metapopulation in the Baltic region remains to be validated through experimental quantification and functional assays. Unfortunately, despite the integration of metagenomic data in our analyses, we were only able to comprehensively investigate the dynamics of the bacterial communities associated with *M. cinxia*, while other microorganisms, including fungi could similarly be affected by climatic variables. Future analyses expanding beyond bacterial communities will be needed to fully assess the impact of climate change on wild host-microbiome-pathogen interactions.

### 4.1 Larvae associate with similar bacteria over a 28-year period

The *M. cinxia* larvae were consistently colonized by similar bacteria across time. The dominant taxa include members of the genera *Pseudomonas* and *Telluria,* and of the family Enterobacteriaceae, broadly consistent with previous work on this host species [33]. Microbiome stability has been reported for diverse herbivores [70–72], and *M. cinxia* larvae could consistently recruit similar bacterial taxa from the predictable source of microbes associated with their host plant and environment [36]. Genomic analyses of the dominant bacterial taxa (i.e., *Pseudomonas*, *Telluria,* and Enterobacteriaceae bacteria), showed that these bacteria encode near-complete pathways for aerobic respiration and substrate transport. These pathways are characteristic of microbes involved in active metabolic exchanges with their host, rather than of transient microbes [73]. Among them, some Enterobacteriaceae bacteria showed a broad repertoire of carbohydrate-active enzymes, including those involved in the degradation of plant cell-wall components [74]. Such bacteria could contribute to the digestion of complex host plant carbohydrates, as previously suggested in the interaction between bacteria *Proteus vulgaris* and *Serratia liquefaciens* and their host, the silk moth *Bombyx mori* [74].

The co-occurrence of multiple Enterobacteriaceae with overlapping carbohydrate-degradation capacity implies a degree of functional redundancy within this clade. Functional redundancy buffers microbial communities against the loss of any single taxon (ref). Previous experiments with laboratory-reared larvae found no effect of gut bacterial composition on larval performance or metabolism, although the microbiota did modulate immune gene expression [34]. The gut microbiota of laboratory larvae, however, differed markedly from the bacterial communities characterized in wild specimens collected for this study. In *M. cinxia*, we would need absolute abundance or transcriptomic data to determine whether the simultaneous loss of several Enterobacteriaceae taxa corresponds to actual losses in the host’s metabolic capacity.

Noticeably, the metabarcoding and the metagenomic approaches did not yield the same data, as some taxa amplified through the metabarcoding approach were absent from the metagenomic data. This difference results from limitations of the sequencing approaches, with the metabarcoding potentially leading to the overrepresentation of some taxa through PCR amplification, including transient diet-associated taxa, whereas metagenomic sequencing possibly more accurately capture the true prevalence of transient and resident microbial taxa.

### 4.2 Effects of three decades of climatic fluctuations

Over the 28-year period studied here, our analyses show that the relative importance of several bacterial genera fluctuated with local temperatures and/or precipitations.

The abundance of the *Telluria* bacteria was positively associated with temperatures, while the abundance of *Pseudomonas* increased with precipitations. Climate-associated changes in species gut microbiota composition have been illustrated for few wild and many lab populations. For instance, *Bacillus* replaced *Pseudomonas* in the sandfly *Lutzomyia longipalpis* when reared under elevated environmental temperatures [75], while a 2°C increase in temperatures resulted in a clear shift in the microbiota in *Aedes aegypti* mosquitoes [76]. Similarly, climate change mediated changes in the diet of the spotted hyenas, leading to a shift in the carnivore’s microbiota from being dominated with *Firmicute* bacteria to being dominated by *Bacteroidales* bacteria, illustrating the plasticity of these carnivores to changes in their foraging habits. In *M. cinxia*, the fluctuations of Telluria and Pseudomonas could be a direct consequence of the effect of the climatic variables on growth of the bacteria in their host gut system. Although *Telluria* and *Pseudomonas* bacteria are both known to grow best under warm temperatures (25-35°C), only the *Pseudomonas* bacteria have been shown to form biofilms when exposed to cooler and wetter conditions [77]. Alternatively, fluctuations in the abundances of these bacteria might be the indirect result of changes in their abundances on the leaf-surface of P. lanceolata in response to changes in the plant’s moisture, chemistry, or physiology [78]. Climate could also slowly modulate the filtering of bacteria in *M. cinxia* by altering the host plant nutritional quality [17] or the host physiology [79].

### 4.3 Impact of the 2018 drought

The drought of 2018 offered a natural setting to test the effect of an extreme climatic event, characteristic of climate change, on the microbiota associated with a wild species. Our comparative analyses of the genomic data from before and after the drought revealed that the most severely affected bacteria belonged to the Enterobacteriaceae family. Bacteria such as *Serratia sp., Pantoea sp.*, *Citrobacter sp.* were prevalent in the gut of *M. cinxia* larvae in the years prior to the drought (e.g.

2011-2012), but all became rare in 2018. A similar pattern was also observed in non-Enterobacteriaceae *Rhabdochlamydia sp*., and *Acinetobacter guillouiae*. Meanwhile, other taxa such as *Hafnia alvei*, *Pseudomonas_E parakoreensis*, and *Aquirickettsiella* sp. showed higher relative abundances in 2018.

We further documented the resilience of the microbiota of *M. cinxia* larvae under natural settings, by highlighting a delayed recovery of the microbiome, which had not been and could not be characterized under previous short-term laboratory conditions. The bacterial taxa depleted by the 2018 drought did not rebound immediately, but returned to pre-drought levels only by 2020, under climatic conditions more typical of the local environment (Figure S14–15; Table S12). Compared to Rosa et al. (2019), who found that moderate drought stress increased gut microbiota heterogeneity in a lab experiment. Our results suggest a directional shift without loss or gain in diversity, rather than stochastic diversification. This difference may reflect the greater severity of the 2018 drought, or other unknown factors present under natural but not laboratory settings [17]. Although laboratory studies in insects have previously demonstrated drastic collapse of bacterial species under heat stress [17, 80], sometimes followed with recovery to pre-stress levels [81], it remains difficult to accurately predict microbiome responses to climate change from controlled experiments alone. As climate models predict accelerated warming, and increasingly variable precipitation regimes in northern Europe, wild species will experience humid winters and more intense summer droughts [82]. Under these conditions, both wet- and drought-associated taxa may be alternately promoted, amplifying temporal fluctuations in wild species associated microbiota composition.

### 4.4 Candidate pathogens

Fluctuations in climatic variables are well-known to affect the prevalence of pathogens [19]. Through functional analyses of *M. cinxia* metagenomes, we found that two MAGs fluctuating with climatic variables, *Enterobacter ludwigii* and *Hafnia alvei*, encoded complete versions of the Na -NADH-ubiquinone oxidoreductase pathway, generally considered a pathogen-specific pathway [83]. Although the bacterium *Enterobacter ludwigii* is non-pathogenic to one of its natural hosts, the moth *Helicoverpa zea* [84], it was shown to induce neurodegeneration in *Drosophila melanogaster* [85]. Similarly, the bacterium *Hafnia alvei* was shown to cause septicemia in honeybees [86]. Although the pathogenic phenotypes of these infections in *M. cinxia* remain to be characterized, our analyses suggest that climatic extremes may favor larval infection with opportunistic pathogens, with *Enterobacter ludwigii* being more prominent in wetter years and *Hafnia alvei* increasing under drier conditions.

## 5. CONCLUSION

The unique long-term data of the butterfly *Melitaea cinxia* metapopulation on the Åland islands reveals complex shifts in this wild species’ associated microbial communities, which experienced both slow and sudden extreme climatic changes. Although many host-bacteria interactions showed some level of resilience to, and delayed recovery from an extreme drought event, bacteria with potentially key functional roles for their butterfly host *M. cinxia* were affected. The predicted accelerating warming and increasing frequency of droughts (or other extreme climatic events) might permanently disrupt the *M. cinxia* microbiome in the future, with consequences for the persistence of the Åland butterfly metapopulation in the Baltic. The continuous survey of this butterfly population will be key to further characterizing the role of the microbiome in this wild species response to future changes.

## Acknowledgements

We would like to thank the members of the ISEE and life-history evolution groups for discussions on the study. Thanks to the field assistants, S. Ikonen, and K. Raveala for contributing to the yearly Åland metapopulation survey in the field. Thanks to N. Verspagen for providing the *M. cinxia* lifecycle figure elements. Thanks to T. Hannunen from the Finnish Institute for Molecular Medicine (FIMM) for supervising the sequencing of the 16S rRNA amplicons. We wish to acknowledge CSC – IT Center for Science, Finland, for computational resources, FIMM and Novogene Europe for sequencing services, and the Molecular Ecology and Systematic laboratory (Mes-lab) at the University of Helsinki for providing access to state-of-the-art molecular facilities.

The study was funded by the Research Council of Finland (grants # 321543 and #355152) to AD, and a HiLIFE Fellowship to AD.

## Data availability statement

Raw sequencing data generated in this study have been deposited in the European Nucleotide Archive (ENA) under BioProject accession number PRJEB112583. The code used to conduct the analyses is available at https://github.com/LinyangSun/M.cinxia-Microbiome.

## Notes

### Competing Interest Statement

The authors have declared no competing interest.

### Summary of Updates

update a new version of mauscript

