## Supplementary material for "The microbiota of a wild butterfly species over three decades of climate change": Supplimentary File

Annex S1: Detailed description of the sample preparation and bioinformatic analyses

1. **DNA extraction, PCR amplification, and sequencing**

Through the years, the DNA from all larval specimens was extracted using slightly different methods [1], following progress in DNA extraction methods. Extraction methods include protocols from the Qiagen extraction kits, NucleoSpin 96 Tissue Core Kit (Macherey-Nagel) and salt method. Each DNA extract was individually labeled, and preserved in the MES lab at the University of Helsinki, using a standardized protocol described in [2].

1.1. 16S rRNA metabarcoding

To investigate variation in the microbiota associated with *M. cinxia*, we followed the protocol described by Minard et al [3]. We amplified the hypervariable V5–V6 region of the bacterial rrs gene using the primers 784F (5′-AGGATTAGATACCCTGGTA) and 1061R (5′-CRRCACGAGCTGACGAC) [4], from 570 larval specimens under sterile conditions. Each specimen underwent duplicate PCR reactions, using 2µl of DNA extract. Duplicates from the same specimen/control were later pooled under sterile conditions [5] before being sent for sequencing at the Finnish Institute for Molecular Medicine Finland (FIMM, Finland). Sequencing was conducted on an Illumina MiSeq v3 platform (Illumina, USA) using both forward and reverse primers. To control for potential contamination during the whole procedure (from manipulation of the specimens to DNA extraction, PCR and sequencing), we included 1 negative control (i.e. sterile water) per 96 wells plate (6 in total).

1.2. Whole Genome Sequencing (WGS) metagenomics

To investigate variation in bacterial functionality across years, we analyzed host whole genome sequence data originally produced for another study (van Dis et al., In Prep) to screen for non-host bacterial reads. A subset of 298 larval samples from the original 512 samples were used in this study. All used samples were collected between 2009 and 2021, with only 43 samples overlapping between the two datasets. The DNA from these 298 specimens was extracted using one of three different methods mentioned above. The final concentrations ranged from 4.76-134 ng/μL (median = 35 ng/μL) as measured by Nanodrop or Qubit HS dsDNA kit, and standardized to 15-30 μL volumes.

Samples were sequenced in three batches on Illumina platforms:

- Batch 1 (N=65,samples from 2009-2010) used BGI standard library preparation and was sequenced on NovaSeq PE150 at median 37X coverage (range: 3-75X).
- Batch 2 (N=111,samples from 2018-2019) and batch 3 (N=336, spanning 2009-2021) used Novogene standard library preparation with 350bp inserts and were sequenced on NovaSeq PE150 and NovaSeq X Plus PE150 respectively, both at median 25X coverage (ranges: 21-38X and 20-50X).

Since both NovaSeq platforms use identical quality scoring, all batches were directly integrated into one dataset. Principal component analysis confirmed no batch effects after sequence processing and filtering.

2. **Bioinformatics**

2.1. 16S rRNA metabarcoding

We first checked the quality of the metabarcoding data using FastQC [6]. As the reverse reads were of relatively lower quality, we only used the data from the forward reads for all subsequent analyses. We then used the QIIME2 platform [7] to conduct all the following data cleaning and analyzing steps. Primer sequences were first removed by using the function “cutadapt trim-single”. The function “quality-filter” was then used to filter out sequence with low quality (<20 PHRED score). We performed denoising using the filtered data through “deblur”. The metabarcoding data was also filtered, classified into unique amplicon sequence variants (ASVs), and annotated using the Greengenes2 plugin [8].

Before conducting any bacterial community analyses, we excluded all ASVs represented by fewer than 10 reads across our entire dataset to minimize noise (i.e., the influence of rare taxa). To control for contamination during sample preparation and PCR, the ASVs found in the negative controls were excluded from specimens processed on the same plate. Similarly, specimens displaying low richness (i.e., fewer than 20 observed ASVs) were excluded to reduce biases potentially introduced by low-quality PCR amplifications, or by sequencing anomalies [9]. The threshold of 20 ASVs was determined based on the normal distribution of ASV numbers across our samples. Finally, we rarefied the remaining data to an even depth of 1,500 reads per specimen. This rarefaction depth was determined based on the normal distribution of the sequencing depth across our samples. Using this threshold, we kept 397 out of the 570 specimens originally selected for this study (69.6%) (Fig. S1).

2.2. WGS metagenomic

To screen the non-host data in the *M. cinxia* WGS data for bacterial genomic material, we first removed all host-derived reads using bwa-mem2 v.2.2.1 [10], and checked the quality of unmapped sequences using BBTools and fastp [11]. High-quality reads from bacteria were subsequently assembled into contigs using MEGAHIT [12]. Binning of the resulting contigs was performed using MetaBAT2 [13] with default parameters, followed by refinement using MAGpurify [14], which incorporates taxonomy, tetranucleotide frequency (TNF), and coverage profiles. This approach allowed us to remove incorrectly clustered contigs. The quality of the refined MAGs was further evaluated using CheckM2 [15]. We retained only the MAGs with completeness above 50% and contamination below 10%. The refined MAGs were then deduplicated by using dRep [16], leading to a final list of 25 high quality MAGs.

Taxonomic annotations were only assigned to those 25 high quality MAGs using GTDB-Tk [17]. All 25 MAGs were first annotated using Prokka [18]. KEGG Orthology (KO) assignments were obtained using KofamScan [19], and pathway completeness was assessed using KEGGdecoder (github link: <https://github.com/speeding-up-science-workshops/KEGGDecoder-heatmap>), and carbohydrate-active enzyme capacity was evaluated using dbCAN [20].

We investigated the relationship between each of the 25 high quality MAGs, by reconstructing their phylogenetic relationship, using the species *Bacteroides thetaiotaomicron* as the outgroup of the phylogenetic tree. This outgroup species is commonly found in vertebrate host’s gut microbial communities and is not detected in invertebrate host [21]. First, all bacteria genomes were scanned against the GTDB reference to identify and classify 120 conserved single-copy marker genes. These markers were then aligned and concatenated into a supermatrix [22]. The concatenated alignment was converted to PHYLIP-PAML format and automatically trimmed with trimAl v1.4 [23] to remove poorly aligned regions. Finally, we inferred a maximum-likelihood phylogeny in iqtree2 [24] using the ModelFinder Plus (-m MFP) module to find and fit the best model with 1000 rapid bootstrap replicates.

6.	*FastQC*. 2015.

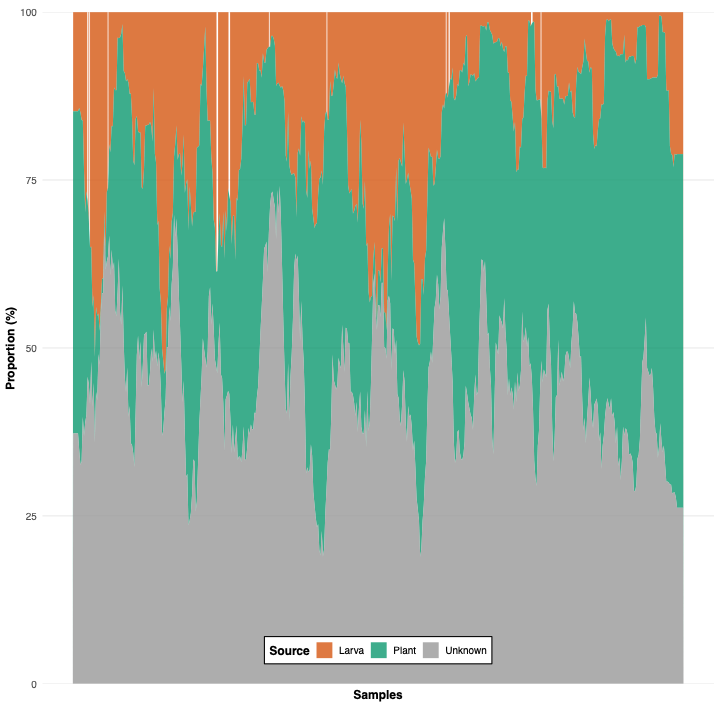

**Figure S1.** Predicted mixing proportions of microbial sources in the 16S rRNA metabarcoding dataset of *M. cinxia* microbiome. Based on a reference database and SourceTracker2 predictions, each bar represents the estimated proportion of larva-associated, plant-derived, and unknown-origin microbes per sample. Plant surface microbes accounted for over one-third (37.21%) of the identified taxa across the dataset.

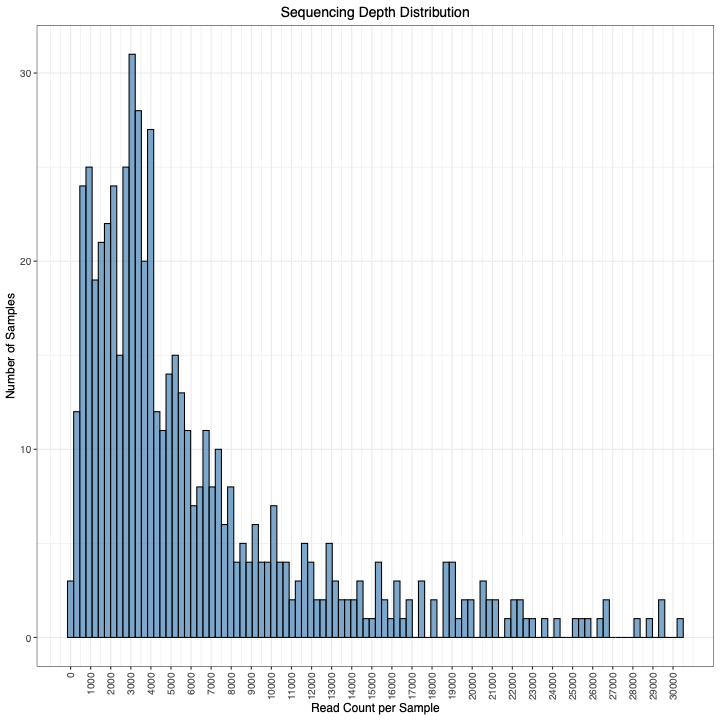

**Figure S2**. Sequencing depth distribution of 16S rRNA metabarcoding data across samples. Distribution of read counts per sample, with the majority of samples yielding between 2,000 and 8,000 reads. The distribution is right-skewed, with a small number of samples exceeding 20,000 reads.

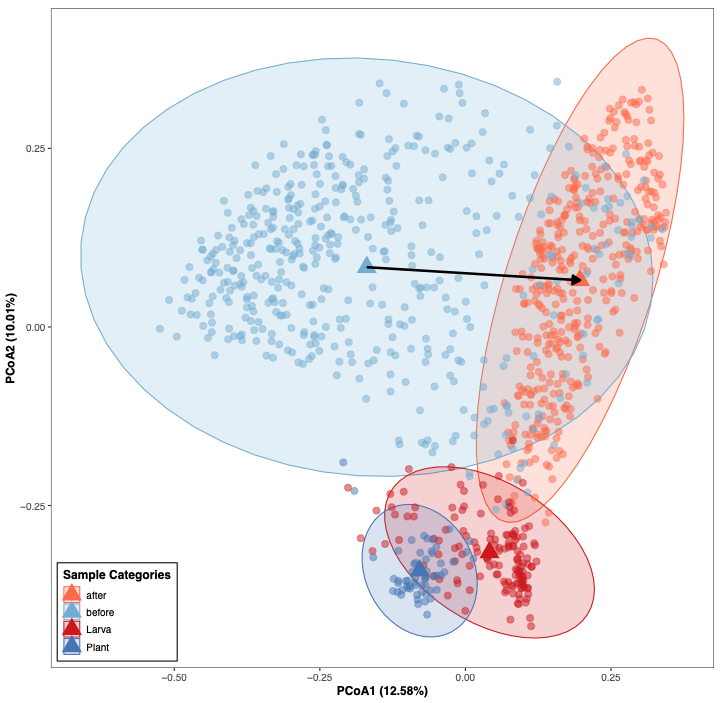

**Figure S3.** Principal coordinates analysis (PCoA) based on Bray–Curtis dissimilarity comparing microbiome community composition before and after removal of plant-derived microbes of 16S rRNA metabarcoding dataset. The first axis (PCoA1: 12.58%) illustrates a clear shift in community composition between pre- and post-removal datasets, alongside reference communities from plant and larva sources.

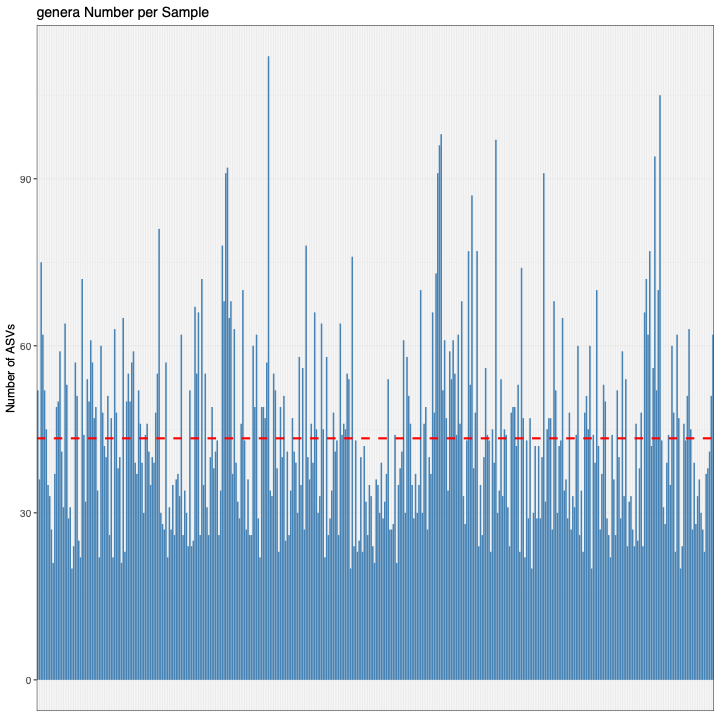

Figure S4. Distribution of the number of microbial genera per sample in 16S rRNA metabarcoding dataset. Across the dataset, individual specimens harbored between 20 and 112 microbial genera, with a median of 41 genera per sample (read dash line).

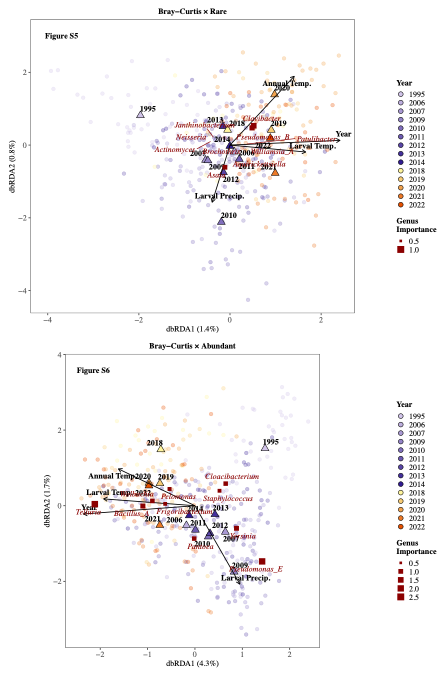

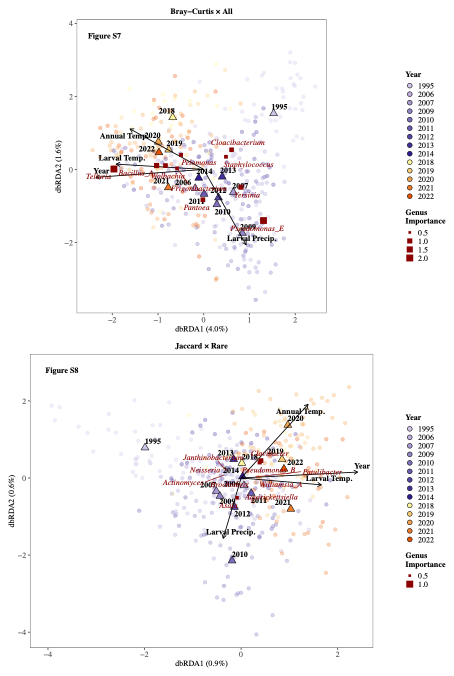

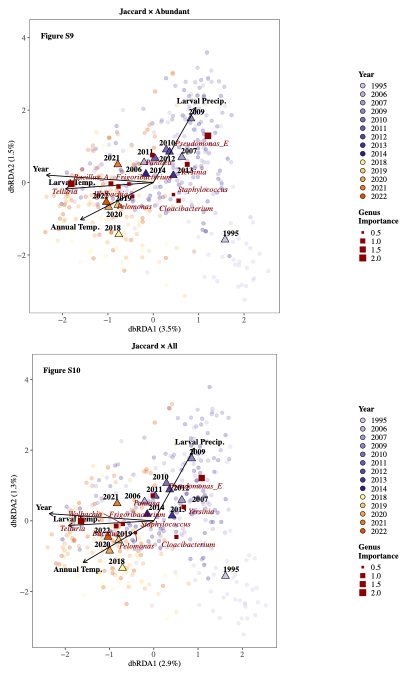

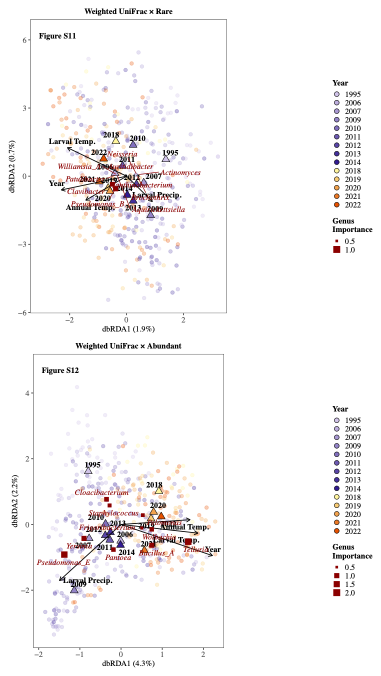

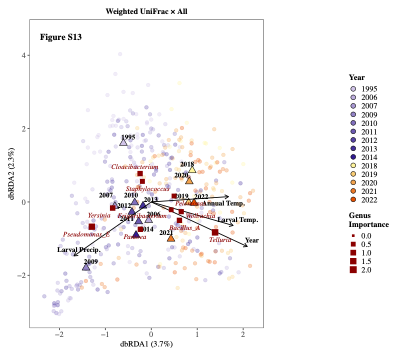

**Figure S5-13.** Distance-based redundancy analysis (dbRDA) ordinations of 16S rRNA metabarcoding bacterial community composition across three OTU subsets (Rare, Abundant, All; columns) and three dissimilarity metrics (Bray-Curtis, Jaccard, Weighted UniFrac; rows). Points represent individual samples colored by sampling year (1995–2022), with shades of purple indicating pre-drought years …. Triangles indicate year centroids and red squares indicate bacterial genera, scaled by genus importance score (0.5–2.0). Arrows represent fitted environmental vectors (Annual Temperature, Larval Temperature, Larval Precipitation, and Year), with direction and length proportional to the strength of association with community composition. Percentages on each axis label indicate the proportion of total variation explained by that constrained axis.

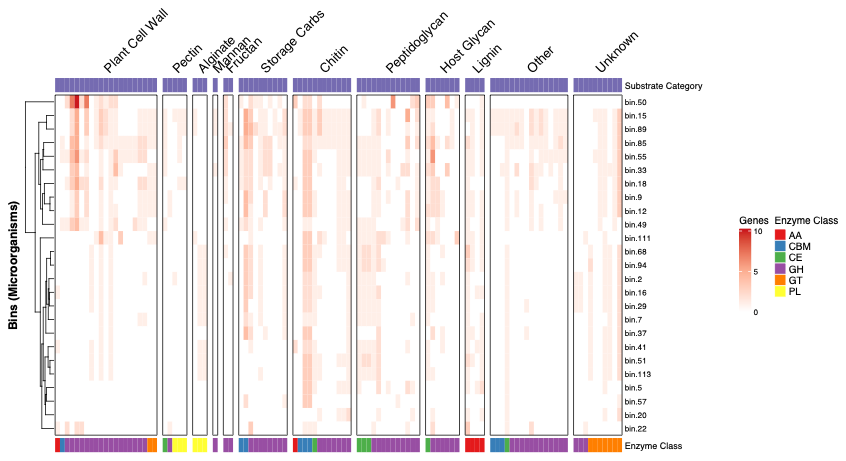

**Figure S14**. Carbohydrate-active enzyme (CAZyme) profiles of metagenome-assembled genomes (MAGs). Heatmap showing the number of genes encoding CAZymes across MAGs, categorized by substrate type (plant cell wall, pectin, alginate, starch/glycogen, storage carbohydrates, chitin, peptidoglycan, host glycan, lignin, and others). Enzyme classes are indicated by color on the left axis (AA, CBM, CE, GH, GT, PL). MAGs and substrate categories are clustered hierarchically.

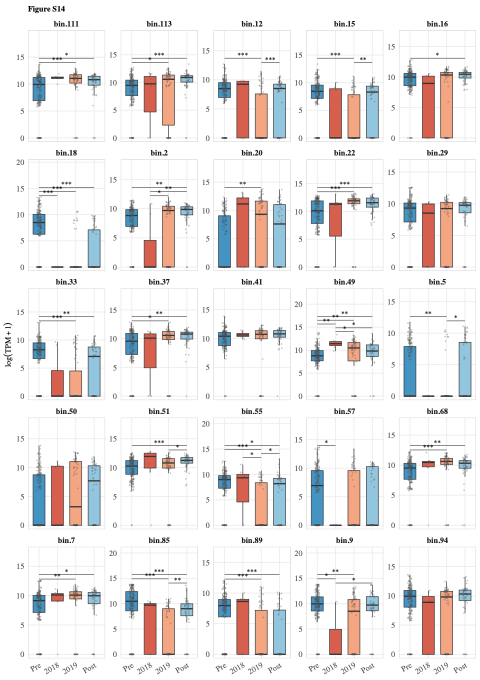

**Figure S14.** Abundance of metagenome-assembled genomes (MAGs) across four periods: Pre-drought (2009–2012), 2018, 2019, and Post-drought (2020–2021). Boxplots show log-transformed TPM (log(TPM + 1)) for each MAG bin across these four periods. Jittered points represent individual samples. Statistical significance was assessed using Kruskal–Wallis tests followed by Dunn's post-hoc pairwise comparisons with Benjamini–Hochberg correction. Significant pairwise differences are indicated by brackets (*P < 0.05, **P < 0.01, ***P < 0.001). Samples with TPM < 300 were set to zero prior to analysis.

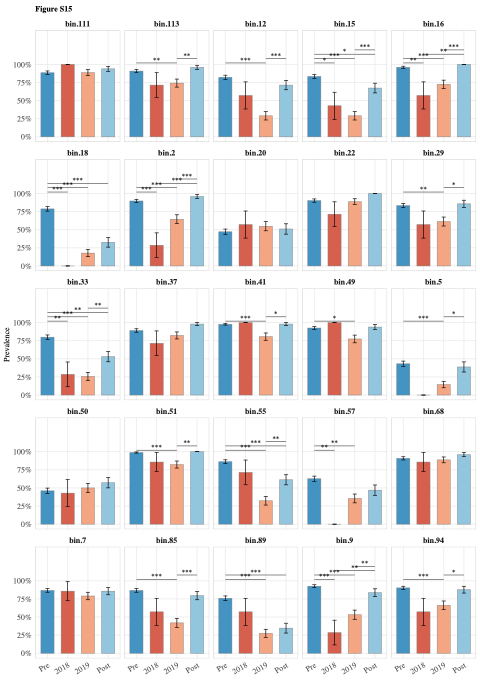

**Figure S15.** Prevalence of metagenome-assembled genomes (MAGs) across four periods: Pre-drought (2009–2012), 2018, 2019, and Post-drought (2020–2021). Bar plots show the proportion of samples in which each MAG bin was detected (TPM ≥ 300) across the four periods. Error bars represent standard errors of the proportion. Statistical significance was assessed using chi-squared tests followed by pairwise Fisher's exact tests with Benjamini–Hochberg correction. Significant pairwise differences are indicated by brackets (*P < 0.05, **P < 0.01, ***P < 0.001).

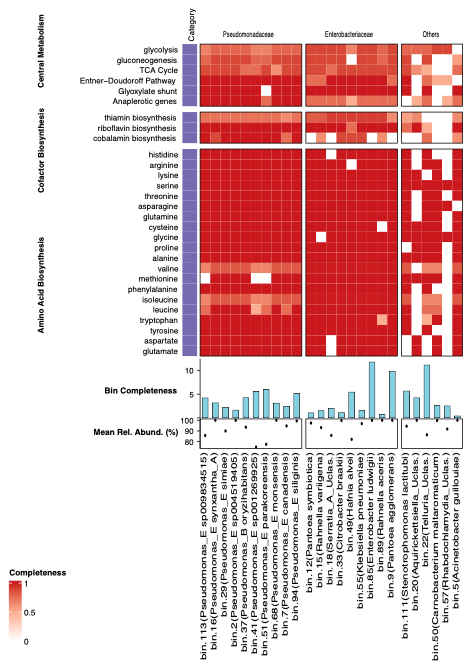

**Figure S16.** Functional potential of 25 high quality metagenome-assembled genomes (MAGs). Completeness of key metabolic pathways across all MAGs, grouped by taxonomic family. Only commonly shared pathways are shown in this figure. Side bar plots indicate mean relative abundance (%) and genome bin completeness (%) for each MAG.
